# Comparison of the two up-to-date sequencing technologies for genome assembly: HiFi reads of Pacbio Sequel II system and ultralong reads of Oxford Nanopore

**DOI:** 10.1101/2020.02.13.948489

**Authors:** Dandan Lang, Shilai Zhang, Pingping Ren, Fan Liang, Zongyi Sun, Guanliang Meng, Yuntao Tan, Jiang Hu, Xiaokang Li, Qihua Lai, Lingling Han, Depeng Wang, Fengyi Hu, Wen Wang, Shanlin Liu

## Abstract

The availability of reference genomes has revolutionized the study of biology. Multiple competing technologies have been developed to improve the quality and robustness of genome assemblies during the last decade. The two widely-used long read sequencing providers – Pacbio (PB) and Oxford Nanopore Technologies (ONT) – have recently updated their platforms: PB enable high throughput HiFi reads with base-level resolution with >99% and ONT generated reads as long as 2 Mb. We applied the two up-to-date platforms to one single rice individual, and then compared the two assemblies to investigate the advantages and limitations of each. The results showed that ONT ultralong reads delivered higher contiguity producing a total of 18 contigs of which 10 were assembled into a single chromosome compared to that of 394 contigs and three chromosome-level contigs for the PB assembly. The ONT ultralong reads also prevented assembly errors caused by long repetitive regions for which we observed a total 44 genes of false redundancies and 10 genes of false losses in the PB assembly leading to over/under-estimations of the gene families in those long repetitive regions. We also noted that the PB HiFi reads generated assemblies with considerably less errors at the level of single nucleotide and small InDels than that of the ONT assembly which generated an average 1.06 errors per Kb assembly and finally engendered 1,475 incorrect gene annotations via altered or truncated protein predictions.

## Main text

The availability of reference genomes has revolutionized the study of biology – many diseases found their causative alleles thanks to the high quality human reference genome [1, 2]; the genomes of agricultural crops have tremendously accelerated our understanding on how artificial selection shaped plant traits and how, in turn, these plant traits may influence species interactions, e.g. phytophagous insects, in agriculture [3, 4]. During the last decade, multiple competing technologies have been developed to improve the quality and robustness of genome assemblies [5–8], enabling genome reference collecting of the tree of life [9–11]. To date, a large number of genomes have been assembled by Third Generation Sequencing (TGS) technologies which can produce individual reads in the range of 10~100 kbs or even longer [12–15]. Although the long-read still has a high error rate, it has been improving owing to the advances in sequencing chemistry and computational tools, e.g. Pacbio (PB) Single-molecule real-time (SMRT) sequencing platform released the Sequel II system of which the updated SMRT cell enabled high throughput HiFi reads using the circular consensus sequencing (CCS) mode to provide base-level resolution with >99% single-molecule read accuracy [16]; while the Oxford Nanopore Technologies (ONT) launched its PromethION platform which can yield > 7 Tb per run and its ultralong sequencing application facilitates the achievement of complete genome - Telomere to Telomere (T2T) - by resolving long and complex repetitive regions for various species including *Homo sapien* [17]. Plenty of species begin to leverage the two cutting-edge sequencing systems; however, almost all chose one single sequencing system, either the PB or the ONT platform, to obtain their reference genomes [15, 18, 19]. Here we present one rice individual (variety 9311) that was sequenced and assembled independently using the two up-to-date systems, and then we compared the two assemblies to investigate the advantages and limitations of each.

Following DNA extraction from the rice sample, we sequenced the two extracts using ONT PromethION and PB Sequel II platforms, respectively. The PromethION generated a total of 92 Gb data with N50 of 41,473 bp and the Sequel II produced a total of 253 Gb data with each molecular fragment being sequenced 14.72 times on average and of an average length of 11,487 bp. We applied multiple software, including Canu1.9 [20], NextDenovo2.0-beta.1 (https://github.com/Nextomics/NextDenovo) and WTDBG2 [21] to assemble the rice genome for the ONT and PB dataset respectively. We selected the optimal assembly for each sequencing platform based on contig N50 and then got both polished (detailed in Supplementary Methods). The ONT assembly showed higher contiguity with a contig number of 18 and an N50 value of ca. 32 Mb in comparison to a contig number of 394 and N50 of 17 Mb for the PB assembly (Figure 1a & Table S1). Ten and three out of the total 12 autosomes were assembled into a single contig in the ONT and PB assembly, respectively. We identified telomers and centromeres for both assemblies (Supplementary Methods) and found that seven of them reached a T2T level assembly with no gaps and no Ns in between (Table S2). A genome completeness assessment using BUSCOv3.1.0 [22] finds both assemblies performed well with the ONT having a tiny improvement (98.62% vs 98.33%, Table S3). We mapped both assemblies to a high-quality rice (R498) genome reference [23] using Minimap2 [24]. Both assemblies showed good collinearity (Figure S1) and the PB assembly contained more gaps in each chromosome compared to that of ONT (Figure 1a).

**Figure 1.**
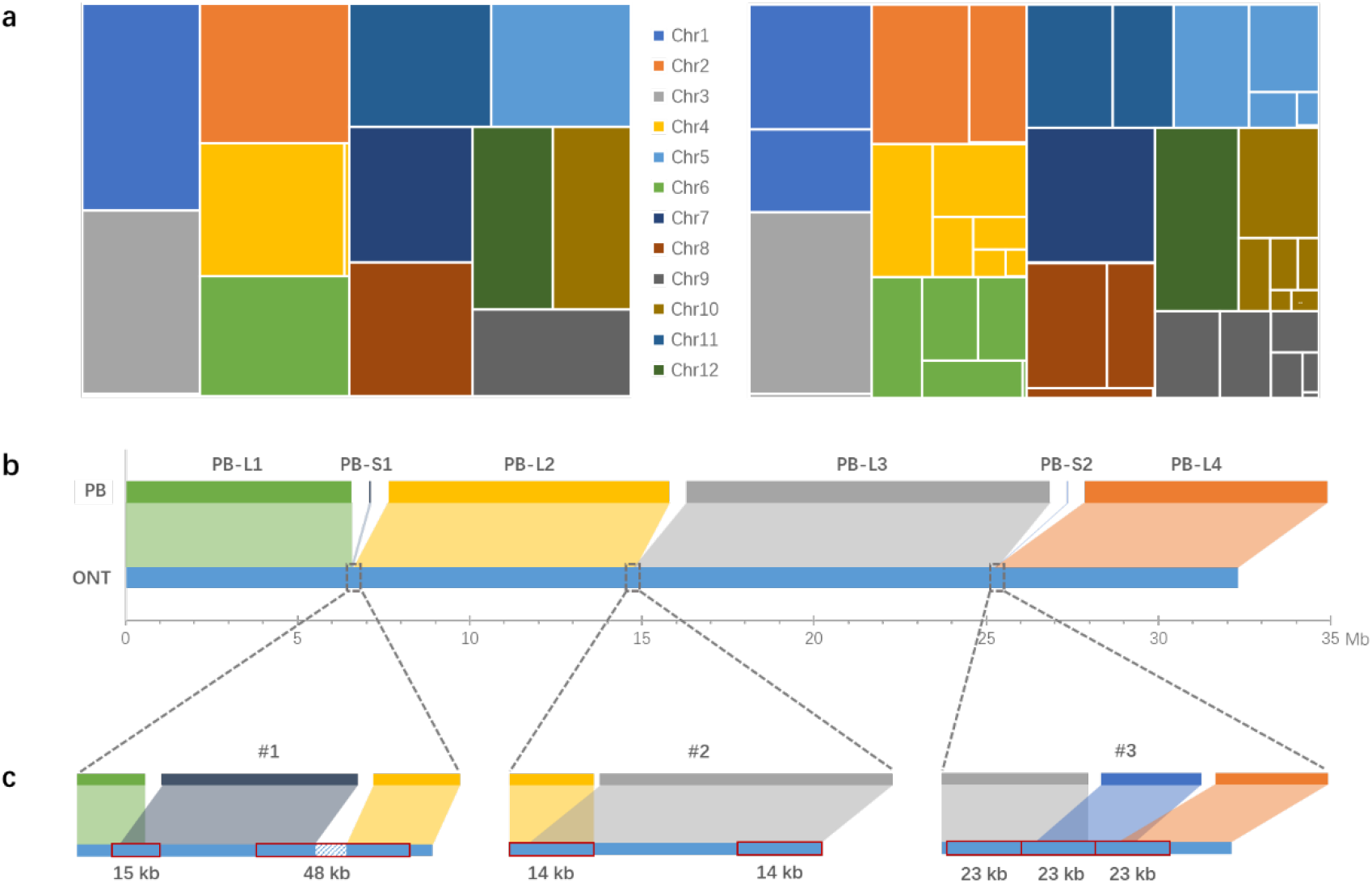
Contiguity of the ONT and PB assemblies. (a) Treemaps for contig length difference between the ONT (left) and PB (right) assembly; (b) The six PB contigs mapped to one ONT contig corresponding to Chr. 6; (c) Details of the three PB gaps. Red rectangles noted the repeat elements.

We then took Chr. 6 where ONT’s one single contig (32,367,127 bp) corresponded to nine contigs (32,476,323 bp) of the PB assembly to investigate and visualize the incongruencies between them. For the nine contigs of PB assembled for the Chr. 6, four reached a length ≥ 6 Mb and five had a length of merely 10-70 kb. We investigated the three gaps where the top four PB contigs (named as PB-L1, PB-L2, PB-L3 and PB-L4 from 5’ to 3’end, respectively) failed to connect (Figure 1b). We mapped the ONT ultralong reads to those gaps and confirmed their correctness through manual inspections by IGV plot [25](Figure S2). The gap #1 between PB-L1 and PB-L2 reached a length of 74,888 bp. One of the short PB contigs (PB-S1, length of 70,208 bp) had an overlap of ~10 kb with the 3’end of PB-L1, thus left the gap #1 a region of 15,722 bp that PB failed to cover (Figure 1c). We further examined the sequences obtained by ONT in and flanking this gap. It showed that the overlapping and the gap regions represented two elements of 15 kb and 48 kb in length that, although have only one copy on Chr. 6, can find their duplications on Chr. 5 (Figure S3). Repetitive elements with such lengths go beyond the typical length generated by PB CCS, therefore the right path can hardly be disentangled from complicated string graphs. The gap #2 between PB-L2 and PB-L3 characterized a region spanning up to 48 kb on the ONT assembly and is flanked by two tandem repeats of 14 kb in length. It was gone through by multiple ONT long reads (Figure S2), so can be successfully connected by the ONT assembly. The last gap between PB-L3 and PB-L4 can be connected by one short PB contig (PB-S2, 25,292 bp), which had 9,469 and 2,621 bp overlaps with 3’end of PB-L3 and 5’end of PB-L4, respectively. And it showed the same case as the gap #2, containing three tandem duplicates of length 23 kb that failed to be connected by PB HiFi reads. We found a total of 107 kb redundancies and 15 kb gaps on Chr. 6 owing to PB’s incorrect assemblies, which corresponded to an excess of 13 annotated genes (Figure 2, Table S4). The genome-wide misassembled regions accumulated to a length of ~ 668 kb (534 kb redundancies and 134 kb gaps), hosting 54 annotated genes (44 redundancies and 10 loss, Table S4). As PB assembly did not generate any single contigs that ONT broke into multiple segments, we cannot find a counter case for comparison.

**Figure 2.**
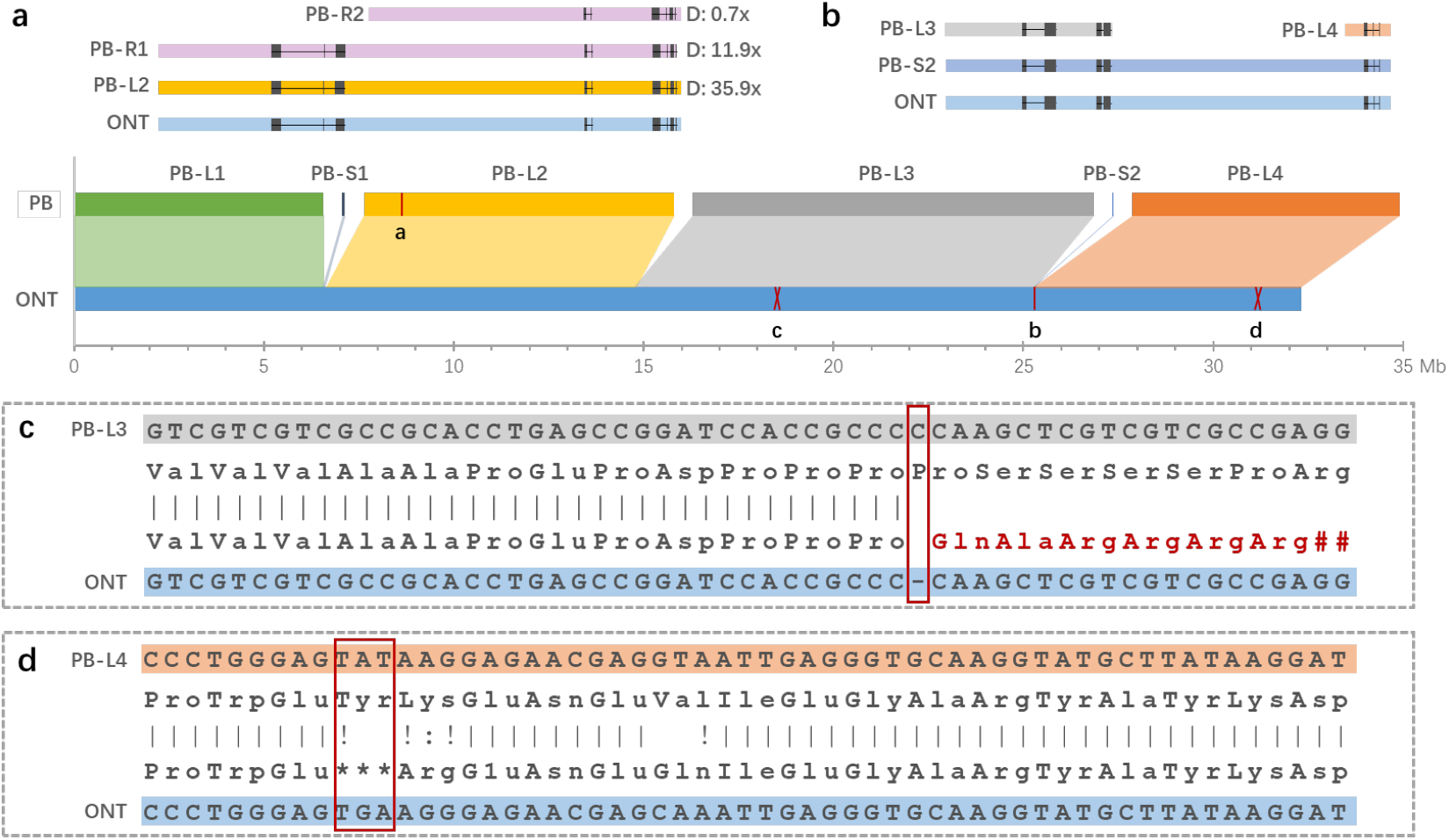
Assembly errors in which genes can be annotated. (a) An example shows gene gains that caused by assembly redundancies, of which the PB-R1 and PB-R2 had a similarity level of 99.67% and 99.51%, respectively, compared to the corresponding region on PB-L2, and “D” abbreviates from depth; (b) The gene redundancies caused by gaps that failed to be correctly connected by the PB assembly; (c) An example shows a 1-base deletion led to frameshift mistake for protein translation; (d) An example shows single base error led to stop codon gain and truncated protein translation.

In addition to those gaps that PB failed to connect, we noticed that there were a bunch of small-scale mismatches between the two assemblies. Firstly, we extracted the reciprocal matches ≥1 M between the two assemblies for comparison using QUAST 5.0.2 [26]. Then, we mapped the PB HiFi reads to both genome assemblies to filter out the innate discrepancies derived from diploid heterozygous states under the assumption that HiFi reads provide high level single base accuracy (Supplementary Methods). It showed that the ONT assembly, although polished using 70× Illumina’s shotgun reads, still contained a large number of small-scale mis-assemblies. In total, we found 210,993 single nucleotide errors and 211,517 InDels (Mean: 1.39 bp, Figure S4) accounting for an average number of 1.06 errors per kb. However, instead of scattering evenly on the assembly, those errors formed into clusters (Figure S5). A further investigation for those regions showed ~ 94% of them have a shotgun read coverage ≤ 5, which explains why the last polishing step failed to fix those errors (Figure S6). About 7.48 % of those errors located on exons and affected ~ 2,415 exons (1,475 genes) to translate correctly to amino acid sequences. We did note that, however, the errors of HiFi reads may be enriched in sequences with particular characteristics, rather than completely random, for example, regions like simple sequence repeats and long homopolymers (Supplementary Methods, Figure S7) which may exacerbate the above error statistics for the ONT assembly.

In conclusion, our study investigated genome assembly qualities between the two up-to-date competing long read sequencing techniques - the PB’s HiFi reads and the ONT’s ultralong reads. It showed both techniques had their own merits with: (1) ONT ultralong reads delivered higher contiguity and prevented false redundancies caused by long repeats, which, in our case of rice genome, assembled 10 out of the 12 autosomes into one single contig, and (2) PB HiFi reads produced fewer errors at the level of single nucleotide and small InDels and obtained more than 1,400 genes that incorrectly annotated in the ONT assembly due to its error prone reads. Therefore, we suggest that further genomic studies, especially genome reference constructions, should leverage both techniques to lessen the impact of assembly errors and subsequent annotation mistakes rooted in each. There is also an urgent demand for improved assembly and error correction algorithms to fulfill this task.

## Availability of data and materials

We have all the data including two genome assemblies and their corresponding raw reads deposited on NCBI under the project ID PRJNA600693.

