## Supplementary information for "Comparison of the two up-to-date sequencing technologies for genome assembly: HiFi reads of Pacbio Sequel II system and ultralong reads of Oxford Nanopore"

### Supplementary Methods

#### Sample preparation and sequencing

We extracted DNA from the rice leaves using SDS method and Q13323kit (QIAGEN) for ONT and PB respectively. We applied SDS method for the ONT platform since ONT ultralong sequencing requires more integrity DNA than general demands of the Third Generation Sequencing. We firstly grinded the leaves carefully with liquid nitrogen and then cleaned cell nucleus with modified buffer HB [1]. After that, the remaining was washed by 1x PBS buffer (pH=7.4, Invitrogen CAT: AM9264) and cell nucleus was lysed by Buffer TLB: 10mM Tris (pH=8.0, Solarbio CAT: T1150), 25mM EDTA, 100mM NaCl (Biosharp CAT: BL542A) and 1.25% SDS (Solarbio CAT: S1015). Next, RNA and protein were removed by RNase A (QIAGEN Lot 8600000227) and protease K (QIAGEN Lot 163024976), respectively. Finally, we eluted the DNA using Buffer EB (QIAGEN CAT: 19086) after phenol/chloroform extracting. Following DNA extraction, two extracts were constructed for ONT and PB libraries and then sequenced using PromethION and PB Sequel II platform, respectively.

We generated a total of 6,100,295 pass reads for the ONT platform with an average quality of 8.99 and N50 of 41,473 bp (Figure S8). PB sequel II platform generated a total of 21,986,306 subreads with each molecular fragment being sequenced 14.72 times on average. It then obtained the HiFi reads from its subreads using the circular consensus sequencing (CCS) mode of pbtools (--min-rq 0.99, --chunk 1/24, -j 4). We removed molecular fragments that were sequenced < 3 times and thus kept a total of 1,494,013 HiFi reads with an average length of 13,363 bp and a genome coverage of 50X.

Additionally, we generated a total of 188,590,034 shotgun reads (~70X) using a strategy of pair end 150 bp (PE 150) on the MGISEQ-2000 platform.

#### Genome assembly

We applied multiple assembly software, including Canu1.9 [2], NextDenovo2.0-beta.1 (https://github.com/Nextomics/NextDenovo) and WTDBG2 [3] to obtain draft genome assemblies for both the ONT and PB platforms (Table S1).

We assembled the PB HiFi reads using CANU1.9 with the parameter -pacbio-hifi, WTDBG2 with -x ccs and NextDenovo2.0-beta.1 with minimap2_options_cns = -x ava-pb -k17 -w17, respectively. For the ONT platform, we applied CANU1.9 with -nanopore-raw, WTDBG2 with -x ont and NextDenovo2.0-beta.1 with correct_option: read_cutoff = 1k, seed_cutoff = 68k. Then, we selected the optimal assembly for each data type - NextDenovo2.0-beta.1 for ONT and Canu1.9 for PB - for the following comparison analyses. After that, we mapped the ONT raw reads and PB HiFi reads onto their corresponding genomes using Minimap2 [4] and conducted genome polishing using RACON [5] for three iterations. Additionally, we applied Medaka, a tool designed for ONT error correction, to polish the ONT assembly. Finally, since HiFi reads of PB platform have already reached an accurate rate of 99% as high as that of shotgun reads of the second generation sequencing platforms, we applied NextPolish1.1.0 [6] to fix the small-scale errors (SNVs and InDels) only for the ONT assembly.

#### Centromere and Telomere sequence identification

We identified centromere and telomere-related sequences for each chromosome-level contig using the RCS2 family repeats and 5‘-AAACCCT-3’ repeats, respectively [7, 8]. For centromeres, we firstly aligned the RCS2 family sequence (AF058902.1) onto the ONT and PB assembly using BWA-mem [9], and then we deemed the regions that contained full units of RCS2 family as centromere repeat regions. For telomeres, we searched for 5‘-AAACCCT-3’ repeats using Tandem Repeats Finder with default parameters [10].

#### Genome annotation

Firstly, we characterized the repeat elements for both PB and ONT genomes using RepeatMasker (http://repeatmasker.org) with a rice repeat library downloaded from the Rice Genome Annotation Project (http://rice.plantbiology.msu.edu/annotation_oryza.shtml). Then, we applied GeMoMa-1.6.1 [11] for protein-coding gene prediction using four related species: *Oryza sativa* ssp. *Indica* (R498, http://www.mbkbase.org/R498/), *Oryza sativa* ssp. *Japonica* (MSU7, http://rice.plantbiology.msu.edu/pub/data/Eukaryotic_Projects/o_sativa/annotation_dbs/pseudomolecules/version_7.0/all.dir/), *Zea mays* (Accession number: GCF_000005005.2) and *Brachypodium distachyon* (Accession number: GCF_000005505.3). After that, we conducted *de novo* gene prediction using Augustus-3.3.1 [12] and combined the two prediction results using EVidenceModeler-1.1.1 [13] with a weight of ABINITIO_PREDICTION (AUGUSTUS) and PROTEIN (GeMoMa) of 1 and 5, respectively.

#### Mismatch and Error detection

*Collinearity:* We aligned both assemblies to a high-quality rice genome (variety R498, Accession ID: GCA_002151415.1) using minimap2 [4] with a parameter setting of -x asm5. Then, we visualized the collinearity between the reference and query genomes using dotPlotly (https://github.com/tpoorten/dotPlotly, -t, -l, -m 30000, -q 1000000).

*Redundancy and gap identification:* We aligned the PB assembly onto the ONT assembly using minimap2 [4] (-x asm5) and kept only the primary hits for each contig. Then, we checked the alignment boundaries of each contig and summarized the redundancies and gaps according to their locations on the ONT assembly. After that, we extracted sequences of the identified gaps and searched for their genome-wide homologous hits.

*Single nucleotide and small Indels:* We aligned PB HiFi reads onto the ONT assembly and then identified SNPs and InDels using GATK4 [14] with filtering parameters: QD < 2.0 || MQ < 40.0 || FS > 60.0 || SOR > 3.0 || MQRankSum < -12.5 || ReadPosRankSum < -8.0 for SNPs and QD < 2.0 || FS > 200.0 || SOR > 10.0 || MQRankSum < -12.5 || ReadPosRankSum < -8.0 for indels. For the incongruencies between ONT and PB assembly, we firstly removed those that were identified to be heterozygous and then tallied the homozygous derived alleles (1/1) that were deemed as ONT errors in the case the derived alleles were identical to that of the PB assembly. We mapped shotgun reads to ONT assembly using BWA-mem [9] and counted the depth of shotgun reads on the ONT errors. For those ONT errors with high shotgun read depth (⩾40), we inspected the alignment between the HiFi read and their corresponding subreads to investigate the potential inaccurate self-corrections during the CCS process. (Figure S7).

#### Comparison of Genes

*Gene loss and redundancies:* For multiple PB assembly contigs that mapped onto the same regions of the ONT assembly, we defined the relatively shorter ones as redundancies if they matched the long contigs with identity > 95% and both their depths were < 40 (Figure 2a). In addition, the gaps (showed in Figure 1) failed to be covered or covered twice by the PB contigs were defined as losses and redundancies, respectively (Figure 2b). And such regions that contained genes contributed the final gene loss and redundancy statistics.

*Incorrectly translation caused by ONT errors:* Firstly, we searched for the ONT errors that located on exons based on gene annotations of both the ONT and PB assembly. For the exon inconsistencies between the two assemblies (present/absent and mismatches), we aligned the amino acid sequences from PB assembly onto the corresponding ONT regions using exonerate [15] (--model protein2genome --refine full -n 1) to investigate how the ONT errors affected gene translation.

#### Supplementary Tables

**Table S1. Assembly quality evaluation**

| | PB HiFi reads | | | ONT ultralong reads | | |
|---|---|---|---|---|---|---|
| | CANU 1.9 (polished) | WDTBG2 | NextDenovo | CANU 1.9 | WDTBG2 | NextDenovo (polished) |
| Total length | 404,883,979 | 351,761,286 | 398,006,903 | 414,548,488 | 678,186,814 | 399,283,841 |
| # of Contigs | 394 | 1,822 | 539 | 961 | 8,982 | 18 |
| N50 | 17,256,392 | 461,009 | 1,663,585 | 3,144,024 | 138,436 | 32,367,127 |
| Average length | 1,027,624 | 193,063 | 738,417 | 431,372 | 75,505 | 22,182,436 |

Note: CANU version 1.9 was updated specially for PB HiFi reads.

**Table S2. The centromeres and telomeres for each chromosome-level contig of ONT and PB assemblies.**

| **ONT contigs** | | | | | |
|---|---|---|---|---|---|
| Contig ID | Centromere position | Long arm length | Short arm length | Telomere repeat number | |
| ctg000009 (Chr.1) | 26,490,151 - 27,493,409 | 26,490,150 | 17,524,811 | 2,450 | - |
| ctg000001 (Chr.2) | 24,270,728 - 24,825,807 | 24,270,727 | 14,198,567 | 2,771 | - |
| ctg000015 (Chr.5) | 12,564,079 - 12,685,655 | 18,233,652 | 12,564,078 | - | - |
| ctg000005 (Chr.6) | 15,722,024 - 15,827,067 | 16,540,060 | 15,722,023 | 2,226 | 2,242 |
| ctg000014 (Chr.7) | 12,648,717 - 13,228,755 | 17,486,620 | 12,648,716 | 1,864 | - |
| ctg000008 (Chr.8) | 12,798,891 - 13,875,118 | 16,574,980 | 12,798,890 | 1,970 | 2,177 |
| ctg000000 (Chr.9) | 2,905,622 - 3,559,788 | 21,497,123 | 2,905,621 | - | - |
| ctg000012 (Chr.10) | 16,285,025 - 16,412,330 | 16,285,024 | 9,681,892 | 1,643 | 2,071 |
| ctg000004 (Chr.11) | 19,266,069 - 20,324,194 | 19,266,068 | 12,803,958 | 1,556 | - |
| ctg000010 (Chr.12) | 11,433,684 - 11,550,742 | 15,397,670 | 11,433,683 | 1,660 | 1,683 |
| **PB contigs** | | | | | |
| Contig ID | Centromere position | Long arm length | Short arm length | Telomere repeat number | |
| tig00000013 (Chr. 3) | 21,058,574 - 21,265,342 | 21,058,573 | 18,168,380 | 3,852 | 3,192 |
| tig00000139 (Chr. 7) | 17,493,629 - 18,180,458 | 17,493,628 | 12,650,984 | 1,718 | 2,642 |
| tig00000277 (Chr. 12) | 15,413,384 - 15,530,454 | 15,413,383 | 11,436,959 | 1,670 | 1,883 |

Note: Telomere repeat number represents the number of 5‘-AAACCCT-3’ repeat with – means fail to find.

**Table S3. Results of genome completeness assessment using BUSCO**

| | PB assembly (%) | ONT assembly (%) |
|---|---|---|
| Complete BUSCOs | 98.33 | 98.62 |
| Complete and single-copy BUSCOs | 96.36 | 96.80 |
| Complete and duplicated BUSCOs | 1.96 | 1.82 |
| Fragmented BUSCOs | 0.29 | 0.15 |
| Missing BUSCOs | 1.38 | 1.24 |
| Embryophyta | 100.00 | 100.00 |

NOTE: PB assembly represents the one assembled using CANU 1.9 and ONT assembly was obtained using NextDenovo.

**Table S4. Gene loss and redundancies of the PB assembly**

| **Gene loss** | | | | |
|---|---|---|---|---|
| ONT_contig_ID | ONT_gene_ID | Start | End | # of loss |
| ctg000002 | ctg000002_G00074 | 999423 | 1034123 | 1 |
| ctg000002 | ctg000002_G00077 | 1062799 | 1069109 | 1 |
| ctg000004 | ctg000004_G01301 | 19428988 | 19446007 | 1 |
| ctg000008 | ctg000008_G01382 | 19347370 | 19353870 | 1 |
| ctg000008 | ctg000008_G02418 | 29556208 | 29684974 | 1 |
| ctg000012 | ctg000012_G00322 | 2969329 | 2970124 | 1 |
| ctg000012 | ctg000012_G00567 | 5045462 | 5046104 | 1 |
| ctg000012 | ctg000012_G00568 | 5049836 | 5052763 | 1 |
| ctg000012 | ctg000012_G00569 | 5053647 | 5056363 | 1 |
| ctg000012 | ctg000012_G00570 | 5056931 | 5058839 | 1 |
| **Gene redundancies** | | | | |
| ONT_contig_ID | ONT_gene_ID | Start | End | # of redundancies |
| ctg000000 | ctg000000_G00597 | 11278491 | 11278830 | 2 |
| ctg000000 | ctg000000_G00598 | 11279619 | 11293649 | 1 |
| ctg000000 | ctg000000_G00081 | 1367697 | 1385458 | 1 |
| ctg000000 | ctg000000_G01574 | 21546713 | 21547382 | 1 |
| ctg000002 | ctg000002_G00074 | 999423 | 1034123 | 1 |
| ctg000002 | ctg000002_G00079 | 1080239 | 1107445 | 1 |

| | | | | |
|---|---|---|---|---|
| ctg000003 | ctg000003_G00812 | 15707253 | 15728302 | 1 |
| ctg000004 | ctg000004_G00097 | 1418133 | 1418904 | 1 |
| ctg000004 | ctg000004_G00098 | 1420869 | 1421634 | 1 |
| ctg000005 | ctg000005_G00255 | 1931660 | 1932002 | 1 |
| ctg000005 | ctg000005_G00256 | 1933158 | 1934211 | 1 |
| ctg000005 | ctg000005_G00257 | 1940317 | 1945807 | 1 |
| ctg000005 | ctg000005_G00258 | 1947015 | 1948068 | 1 |
| ctg000005 | ctg000005_G00932 | 7747015 | 7751198 | 1 |
| ctg000005 | ctg000005_G00933 | 7762171 | 7762622 | 2 |
| ctg000005 | ctg000005_G00934 | 7766283 | 7780469 | 2 |
| ctg000005 | ctg000005_G01464 | 14805155 | 14815093 | 1 |
| ctg000005 | ctg000005_G02124 | 25314455 | 25316369 | 1 |
| ctg000005 | ctg000005_G02125 | 25318653 | 25322992 | 1 |
| ctg000005 | ctg000005_G02126 | 25333867 | 25334733 | 1 |
| ctg000006 | ctg000006_G03206 | 33368857 | 33372453 | 1 |
| ctg000006 | ctg000006_G03207 | 33373708 | 33375291 | 1 |
| ctg000006 | ctg000006_G03208 | 33380579 | 33384195 | 1 |
| ctg000006 | ctg000006_G03209 | 33384930 | 33385787 | 1 |
| ctg000006 | ctg000006_G03210 | 33386180 | 33386905 | 1 |
| ctg000008 | ctg000008_G01382 | 19347370 | 19353870 | 1 |
| ctg000008 | ctg000008_G01383 | 19356150 | 19363873 | 1 |
| ctg000008 | ctg000008_G01252 | 17263321 | 17263702 | 1 |
| ctg000008 | ctg000008_G01253 | 17263970 | 17264438 | 1 |
| ctg000008 | ctg000008_G02418 | 29556208 | 29684974 | 1 |
| ctg000012 | ctg000012_G00286 | 2690576 | 2700335 | 1 |
| ctg000012 | ctg000012_G00287 | 2703559 | 2704736 | 1 |
| ctg000012 | ctg000012_G00989 | 9408612 | 9410982 | 1 |
| ctg000012 | ctg000012_G00990 | 9411510 | 9413516 | 1 |
| ctg000012 | ctg000012_G00991 | 9416266 | 9418167 | 1 |
| ctg000012 | ctg000012_G00992 | 9419388 | 9424819 | 1 |
| ctg000015 | ctg000015_G00162 | 1369836 | 1372125 | 2 |
| ctg000015 | ctg000015_G00163 | 1373588 | 1377298 | 2 |
| ctg000015 | ctg000015_G00008 | 83886 | 101996 | 1 |

Note: the start and end positions corresponded to the coordinates on the ONT assembly.

#### Supplementary Figures

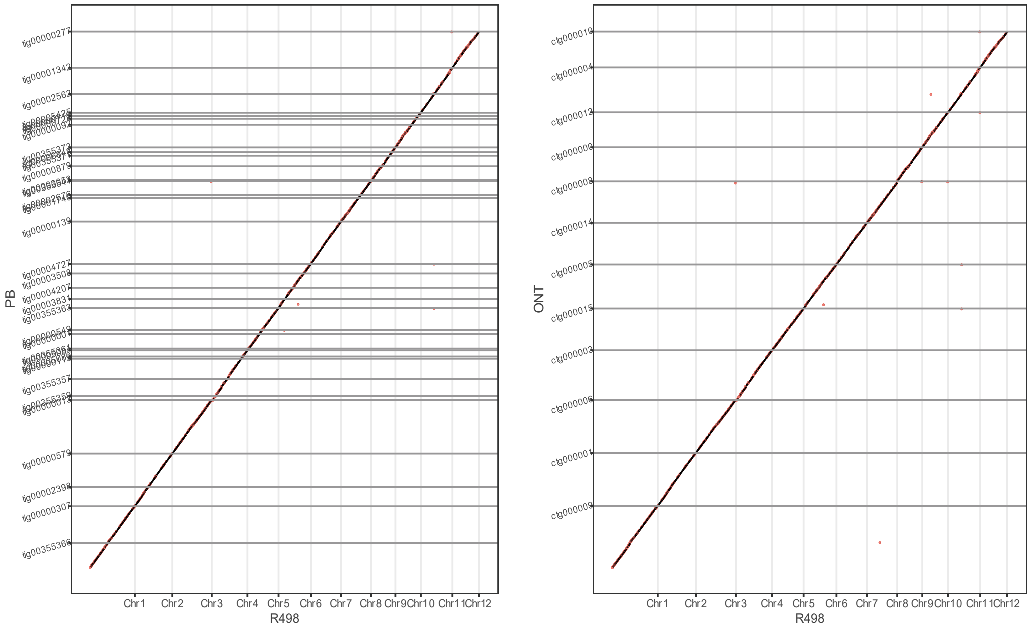


**Figure S1. Collinearity between genome assembly of rice R498 and that of the PB (left) and ONT (right).** Note: It only shows alignments ⩾ 30 kb and query sequences ⩾ 1 Mb.

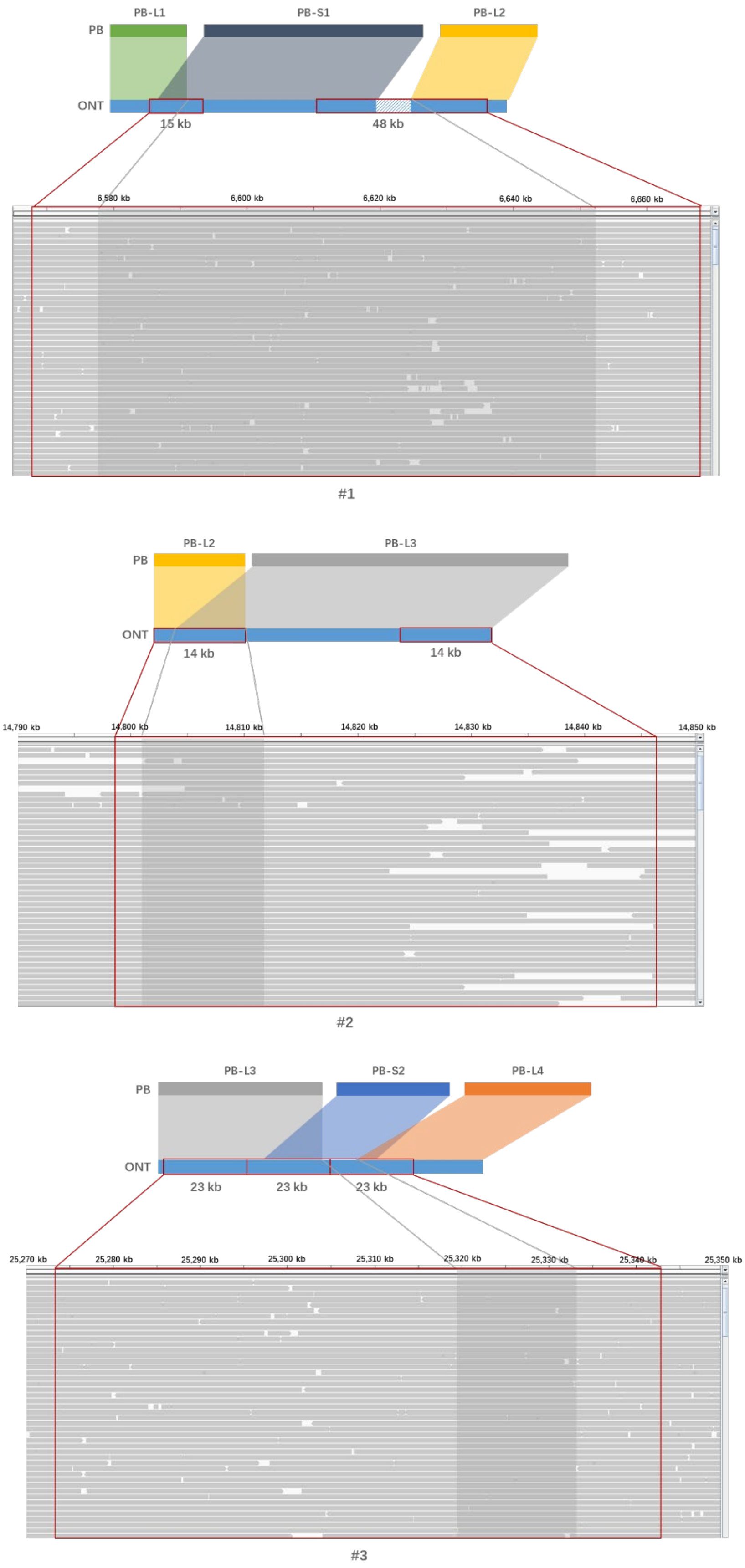


**Figure S2. IGV plots of the three PB gaps on Chr. 6.** Gray shadows represent gap regions of the PB assembly. Red rectangles represent the repeat elements.

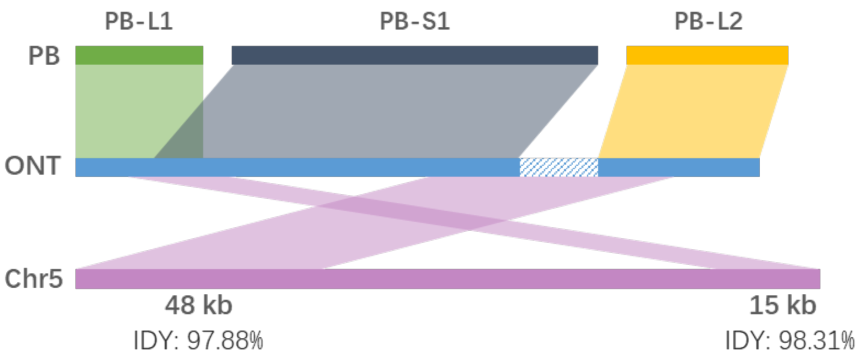


**Figure S3. Details of PB gap #1.** The two repetitive regions matched to another ONT assembly contig with high identities. IDY means similarity identities between each other.

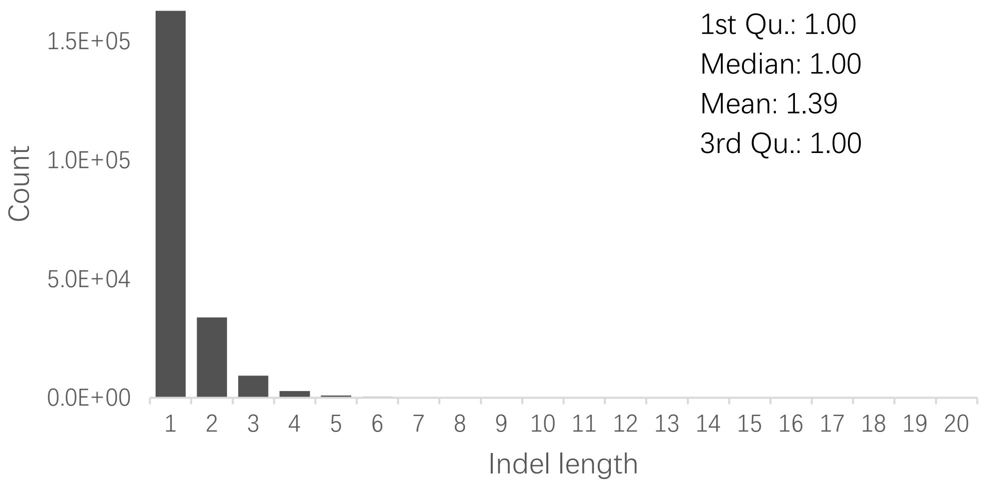


**Figure S4. The length distribution of the ONT InDel errors.** Note that InDels of length > 20 bp had a total count of 216 and did not show here.

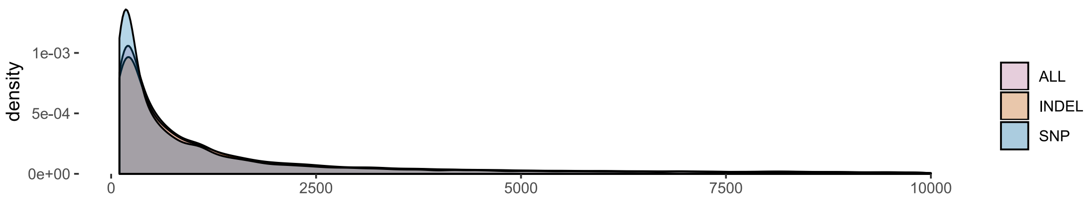


**Figure S5. Distances between adjacent ONT errors.** Those errors tend to cluster in the same region rather than distribute randomly and evenly on the genome, since the distances should have a peak around 1,000 bp for an average error rate of 1.06 per kb in the case of random distribution.

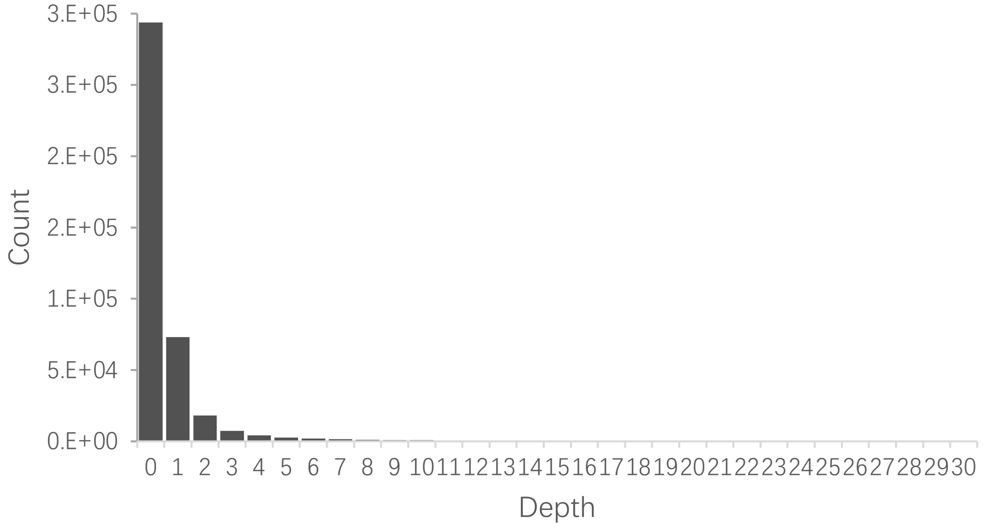


**Figure S6. Depth of Illumina's shotgun reads for the errors of the ONT assembly.** Note that read depth > 30 had a total count of 10,294 (2.44% of total) and did not show here.

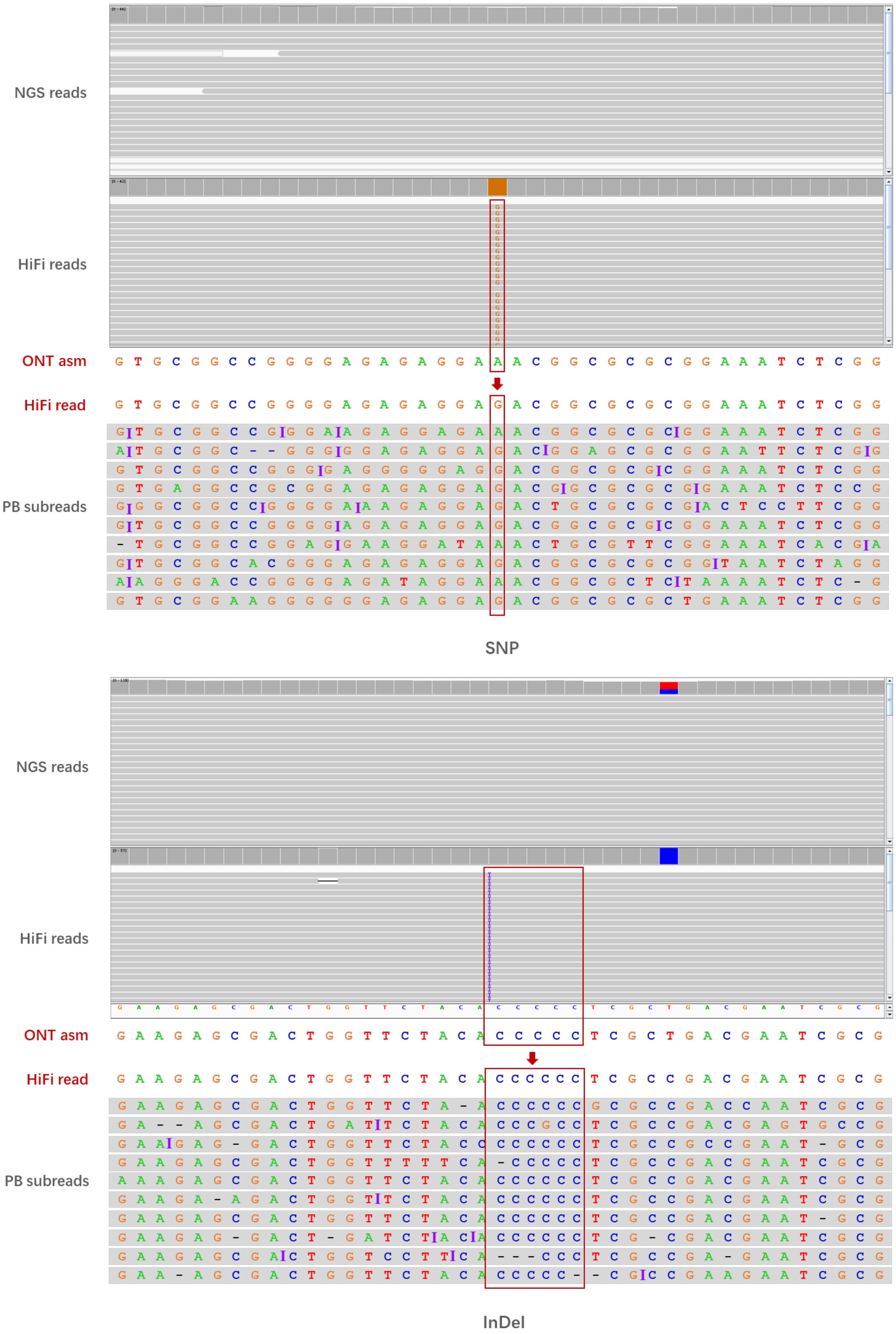


**Figure S7. Two examples (one SNP and one InDel) that showed the mismatches between the ONT and PB assemblies which were well covered by shotgun reads and thus could be errors on HiFi reads generated during the CCS progress.**

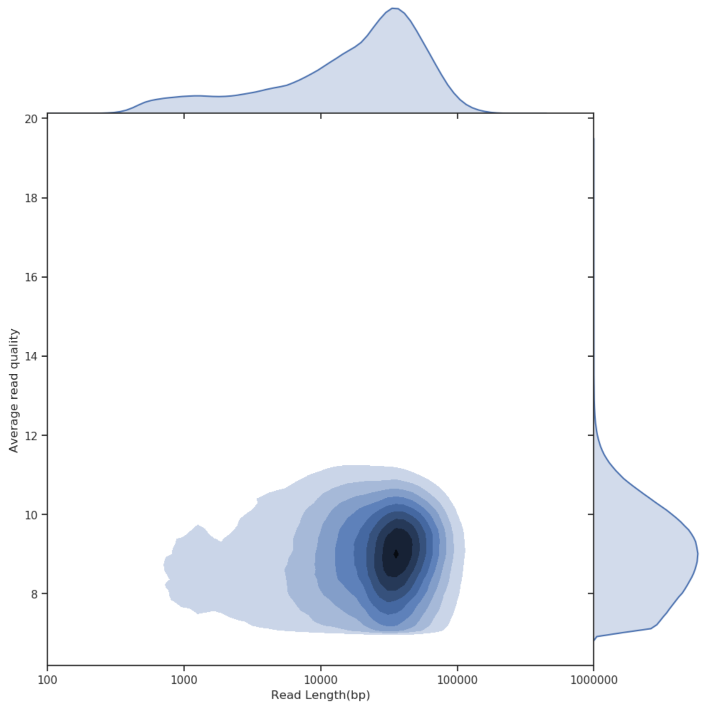


**Figure S8. Quality and length of ONT ultralong reads.** It only showed passed reads with quality ⩾ 7.
